# A Con Artist: Phenylphenoxybenzamide is not a Glycosyltransferase Inhibitor

**DOI:** 10.1101/292912

**Authors:** Gjalt G. Wybenga, Wei-Shen Wu

**Affiliations:** Genomics Research Center, Academia Sinica, 128 Academia Road, Section 2, Nankang, Taipei, 115, Taiwan; Graduate Institute of Life Sciences, National Defense Medical Center, 161 Minquan E. Road, Section 6, Neihu, Taipei 114, Taiwan

**Keywords:** crystallography, enzyme assay design and paper writing (GGW), DNA manipulations, enzyme assay co-design and execution (WSW).

## Abstract

To combat bacterial resistance against antibiotics, glycosyltransferase inhibiting molecules, which block the synthesis of the pre-cursor of the bacterial cell wall, need to be discovered and developed. In this study, we demonstrate that phenylphenoxybenzamide, a salicylanilide, is not a glycosyltransferase inhibiting molecule, despite claims in literature to the contrary, and through our work show that glycosyltransferase construct choice and detergent choice are crucial parameters to consider when designing glycosyltransferase assays that aim to discover and develop molecules that inhibit these types of enzymes.

## Introduction

The emergence of bacterial resistance to common antibiotics [1] has prompted initiatives to search for new antibiotics against existing antibiotic targets, for example, enzymes that synthesize peptidoglycan. Peptidoglycan provides mechanical strength to Gram-positive and Gram-negative bacteria, which allows these bacteria to withstand osmosis-induced turgor pressure caused by ionic strength fluctuations in the bacterium’s environment [2]. Peptidoglycan is synthesized from GlcNAc-MurNAc-pentapeptide-diphosphate-undecaprenyl (lipid II) [3, 4] by class A penicillin binding proteins (PBPs) [5]. The N-terminal domain of a PBP encodes a glycosyltransferase (GT). This GT catalyzes a cation cofactor-dependent reaction that results in the formation of a β-1,4 covalent bond between the C4 O-atom of the GlcNAc sugar of lipid II (bound in the acceptor pocket of the active site of the GT) and the C1-atom of the MurNAc sugar of lipid II (bound in the donor pocket). The product subsequently translates through the active site of the GT, after which another lipid II molecule binds the now vacant acceptor pocket. Consecutive reactions result in a lipid II polymer [6, 7], which during the GT-catalyzed reaction, is integrated into pre-existing peptidoglycan by the C-terminal domain of the PBP that catalyzes a transpeptidase (TP) reaction [5]. While PBPs build up and strengthen the peptidoglycan layer that surrounds a bacterium, lytic transglycosylases weaken the peptidoglycan layer by cleaving the β-1,4 covalent bond created by PBPs [9]. Compounds that bind and inhibit the reactions catalyzed by PBPs will thus negatively affect the strength of the bacterial cell wall and are therefore sought after since they can be potentially used as antibiotics. β-lactam antibiotics inhibit the TP reaction catalyzed by PBPs, however bacteria have developed resistance to these types of antibiotics, rendering them less effective in treating bacterial infections [9, 10]. Therefore attention has shifted towards molecules that inhibit the GT-catalyzed reaction by PBPs. Examples are moenomycins and moenomycin analogues [11], monosaccharides ACL20215 and ACL20964 [12], and lipid II substrate analogues [13, 14]. More recently, non-saccharide molecules, such as salicylanilides [15], Albofungin (a xanthone) and TAN1532B (a benzo[*a*]tetracene) were discovered to inhibit the GT-catalyzed reaction [16]. In this research we employ a monoglycosyltransferase from *S. aureus* (*Sa*MGT), a class A PBP without a TP domain (CAZy: GT51), to reveal that phenylphenoxybenzamide (ppb, fig. 1 (**3**), a salicylanilide) is not a GT inhibitor, and in the process establish GT assay design principles that will facilitate GT inhibitor identification and development.

**Fig. 1:**
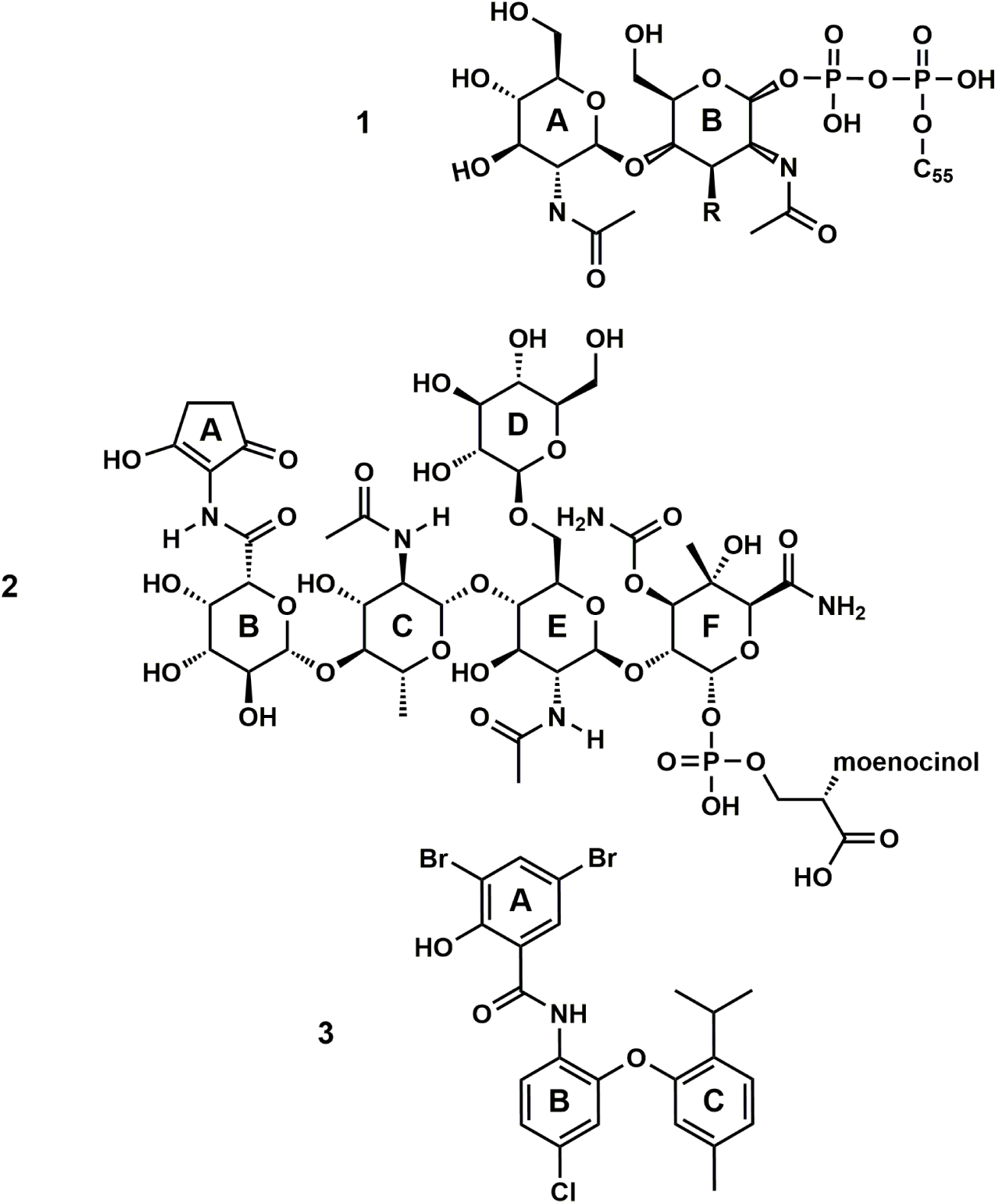
The chemical structures of NBD-lipid II (**1**) with ‘R’ representing a NBD-labeled lactoyl-pentapeptide (AEKAA) as shown in fig. 9B of Cheng *et al*. [15]. Moenomycin A (**2**), a lipid IV product analogue. 3,5-dibromo-N-[4-chloro-2-(5-methyl-2-propan-2-ylphenoxy)phenyl]-2-hydroxybenzamide (ppb, **3**, logP 10.3), a salicylanilide (Cheng *et al*. [15], table 12, compound 42-31). Acquired from Vitas-M laboratory. Figure made with ChemDraw.

## Materials and Methods

### Transformation and Expression of Δ28-269 and Δ68-269 SaMGT

40 ng of DNA (pET15b encoding truncated *Sa*MGT with an N-(Δ28-269, table 1 (2)) or C-terminal linker (Δ68-269, table 1 (3)) containing a His6-tag (confirmed by sequencing)) was added to a vial containing 100 μl thawed *E. coli* Bl21(DE3) competent cells (RBC Biosciences) kept at 4 ºC. The suspension was mixed, transferred to a water bath for transformation (42 ºC, 1 min), returned to ice, and finally pipetted onto a TB agar plate containing 100 μg/ml carbenicillin. The plate was incubated overnight at 37 ºC and bacterial colonies used to inoculate a 100 ml TB pre-culture (100 μg/ml carbenicillin, same temperature, 180 rpm). After 17 h, 5 ml pre-culture was used to inoculate 1 L pre-heated TB medium (50 μg/ml carbenicillin, added prior to inoculation, 37 ºC, 180 rpm) and at OD 0.5-0.6 (reached within 3 h) 0.36 g of IPTG powder was added to induce expression of Δ28-269 (2MGT, Table 1) or Δ68-269 *Sa*MGT (6MGT). After 3 h the medium was collected and centrifuged to harvest the *E. coli* cells, after which the cell pellet was collected, frozen (-20 ºC) and stored until further use.

**Table 1:** An overview of *Sa*MGT constructs used in different studies. 1) *Sa*MGT. UniProtKB Q99T05. Shown for reference purposes. 2, 3) This study. The underlined amino acids were introduced after thrombin cleavage of a thrombin-cleavable linker containing a poly-histidine tag to facilitate nickel affinity chromatography purification. 4) The *Sa*MGT construct used to elucidate the 3D structure of TM-less Δ68-268 *Sa*MGT (68MGT) with bound moeA (PDB code 3HZS). 5, 6) *Sa*MGT constructs used by Wang *et al*. [35] and Terrak *et al*. [36]. Both retain an N-terminal linker containing a poly-histidine tag (H^10^, H^6^, respectively). *The primer translates to R269.

| Nr. | Construct | N-terminus | C-terminus | total number of amino acids | MW (kDa) |
| --- | --- | --- | --- | --- | --- |
| 1 | *Sa*MGT | MKRSDRYSNS | QQAMSQLNR | 269 | 31.459 |
| 2 | Δ28-269 | <u>GSH</u>MQ<sub>28</sub>PVGKP | YQQAMSQLNR<sub>269</sub> | 246 | 28.388 |
| 3 | Δ68-269 | MD<sub>68</sub>NVDELRKI | MSQLNR<sub>269</sub><u>LVPR</u> | 207 | 23.996 |
| 4 | Δ68-268 | <u>NLYFQGH</u>MD<sub>68</sub>N | QYQQAMSQLN<sub>268</sub> | 209 | 24.234 |
| 5 | Δ68-269 | MGH<sup>10</sup>SSGHIEG | YQQAMSQLNR<sub>269</sub> | 224 | 26.052 |
| 6 | Δ68-268* | MGSSH<sup>6</sup>SSGLV | YQQAMSQLNR<sub>269</sub> | 223 | 25.695 |

### Purification of 2MGT

To each gram of frozen cell pellet 10 μl 1 mg/ml DNase I (Sigma) and 20 μl 50 mg/ml lysozyme were added, then 30 ml buffer A (20 mM Tris-HCl pH 8, 200 mM NaCl) and one cOmplete EDTA-free protease inhibitor tablet (Sigma). After thawing, and homogenization by vortexing, the cell solution was passed thrice through a microfluidizer device. The volume of the flow-through was then adjusted to 50 ml with buffer A, n-Decyl-β-D-Maltopyranoside (DM, Anatrace) powder was added (483 mg, 20 mM), incubated (2 h, RT, Intelli Mixer, 4 rpm) and centrifuged (Eppendorf 5810R, rotor F-34-6-38, 10.000 rpm, 10 min) to remove non-solubilized material. The supernatant was decanted, centrifuged once more and loaded onto a 1 ml HisTrap HP column with an ÄKTA FPLC pre-equilibrated with buffer B (20 mM Tris-HCl pH 8, 200 mM NaCl, 50 mM imidazole pH 8 and 2 mM DM). The column was washed with buffer B (15 column volumes), disconnected from the FPLC system, and loaded with 1.1 ml buffer B mixed with 20 μl thrombin (1 unit thrombin per μl 1X PBS) and incubated overnight (23 ºC) to release bound 2MGT. After 20 h the column was reconnected to the FPLC system, washed with buffer B, the flow-through collected, concentrated (Amicon, 10.000 MWCO) and further purified with a Superdex 200 10/300 GL column (GE Healthcare) equilibrated in 200 mM NaCl, 2 mM DM and 20 mM Tris-HCl pH 8 (buffers C) or Na HEPES pH 8 (buffer C’). The purified protein (Figs. S1 & S2) was collected, and either used for crystallization experiments after concentration (buffer C), or frozen (-20 ºC) and after thawing used for enzyme assays (buffer C’).

### Crystallization of 2MGT

For crystallization experiments, purified 2MGT, of which the protein concentration was determined with the Bradford method [17], was mixed with buffer C and bicelle solution resulting in a 15 mg/ml protein, 3 % wt/vol bicelle solution, mixture. A mosquito crystallization robot (TTP Labtech) was employed to set up sitting drop vapor diffusion plates (MRC 2, Hampton) to screen for crystal growth in drops consisting of 0.15 μl protein/bicelle solution and 0.15 μl well solution. Each well (60 μl) of the sitting drop plate contained 1 of 96 different combinations of 100 mM MgCl_2_ (Hampton), 0.1 M Na HEPES pH 8 (Hampton), and 12-34 % wt/vol PEG200-1500 (Hampton, Sigma, Fluka). The plates were sealed with clear sealing tape (Hampton) and immediately transferred to the cold room (4 ºC).

### Preparation of bicelle solution

To make bicelle solution [18] 5 mg CHAPS detergent (GE), 15 mg DMPC (Avanti Polar Lipids) and 100 μl ultra-pure water (Milli-Q) were mixed, heated to 42 ºC (10 min), vortexed, put on ice (10 min), vortexed again, and these steps repeated until all solid material was dissolved. The solution was kept on ice for use in crystallization experiments, or was frozen (-20 ºC).

### Diffraction data processing

The diffraction data collected from a 2MGT crystal were indexed with Mosflm (space group P1) [19], scaled and merged with Scala, after which a self-rotation function was calculated with Molrep. This revealed the presence of three perpendicular 2-fold axes. The data were indexed, scaled and merged again (P222) and the presence of two screw axes established along **a** and **c** (confirmed with Pointless). To generate P2_1_2_1_2 from P222 Reindex was applied with operator h=l, k=h, l=k. The data were then truncated (ellipsoidally) and scaled (anisotropically) using the diffraction anisotropy server (https://services.mbi.ucla.edu/anisoscale/) [20] and reflections assigned to R_free_ (5.6 %) with Uniquefy. The solvent content was determined with Matthews [21] and for phasing Phaser (for molecular replacement) [22] was employed with an input model based upon the model associated with PDB code 3VMQ [23] (without the TM-domain). The model was subsequently refined with Refmac [24], and manipulated with Coot [25]. Scala, Molrep, Pointless, Reindex, and Uniquefy were accessed via the CCP4 software suite [26]. Refinement statistics for the model are reported in Table S1.

### Construction, transformation, expression and purification of transmembrane domain-less 6MGT

The transmembrane (TM) domain of 2MGT was identified with Phobius [27] (see Results) and two primers 5’ CATGCCATGGATAATGTGGATGAACTAAG -3’ (forward) and 5’ CGCGGATCCTCATCAGTGATGATGATGATGATGGCTGCTGCCGCTGCCGCGCGGCACCAGACGATTTAATTGTGACATAGCC -3’ (reverse) designed to construct 6MGT from the 2MGT encoding pET15b plasmid. After PCR the DNA product was purified, restricted with NcoI and BamHI, ligated into a pET15b vector and sequenced. After transformation of *E. coli* Bl21(DE3) cells with the 6MGT encoding plasmid, 6MGT was expressed and purified as described (2MGT) although the last purification step utilized a Superdex75 10/300 GL column equilibrated in buffer C’ without DM (Figs. S1&S2).

### Enzyme assay design

To test whether purified MGT (2MGT or 6MGT) was able to polymerize NDB-lipid II (Fig. 1 (**1**)), NBD-lipid II (lipid II) was pipetted into an eppendorf tube from a stock solution dissolved in MeOH. The tube was transferred into a fume-hood, and the MeOH allowed to evaporate. A buffer, a catalyst, a detergent and/or organic solvents were added followed by purified enzyme. This resulted in an enzyme assay volume of 20 μl, an enzyme concentration of 0.2 μM and a lipid II concentration of 4 μM. The eppendorf tube was then, without delay, transferred into an eppendorf thermomixer R (1400 rpm), and the enzymatic reaction (after time) stopped by the addition of 1 μM moenomyin A (moeA, fig. 1 (**2**), [28]). Next, 200 μg lysozyme was added to convert lipid II polymer into NBD-GlcNAc-MurNAc-dissacharide (room temperature, 2 h) and the mixture analyzed as described by Wu *et al*. [16]. To test the inhibition of MGT by moeA, moeA was added to the enzyme assay solution prior to adding purified enzyme. To test whether MGT was inhibited by ppb (Fig. 1 (**3**)) or TAN1532B (Fig. S4 (**4**), 2MGT only) either compound (dissolved in MeOH) was added to an eppendorf tube together with lipid II, then placed in a fume-hood, and after evaporation of MeOH, used to assay enzymatic activity (Figs. 4, 5 and 6).

**Fig. 2:**
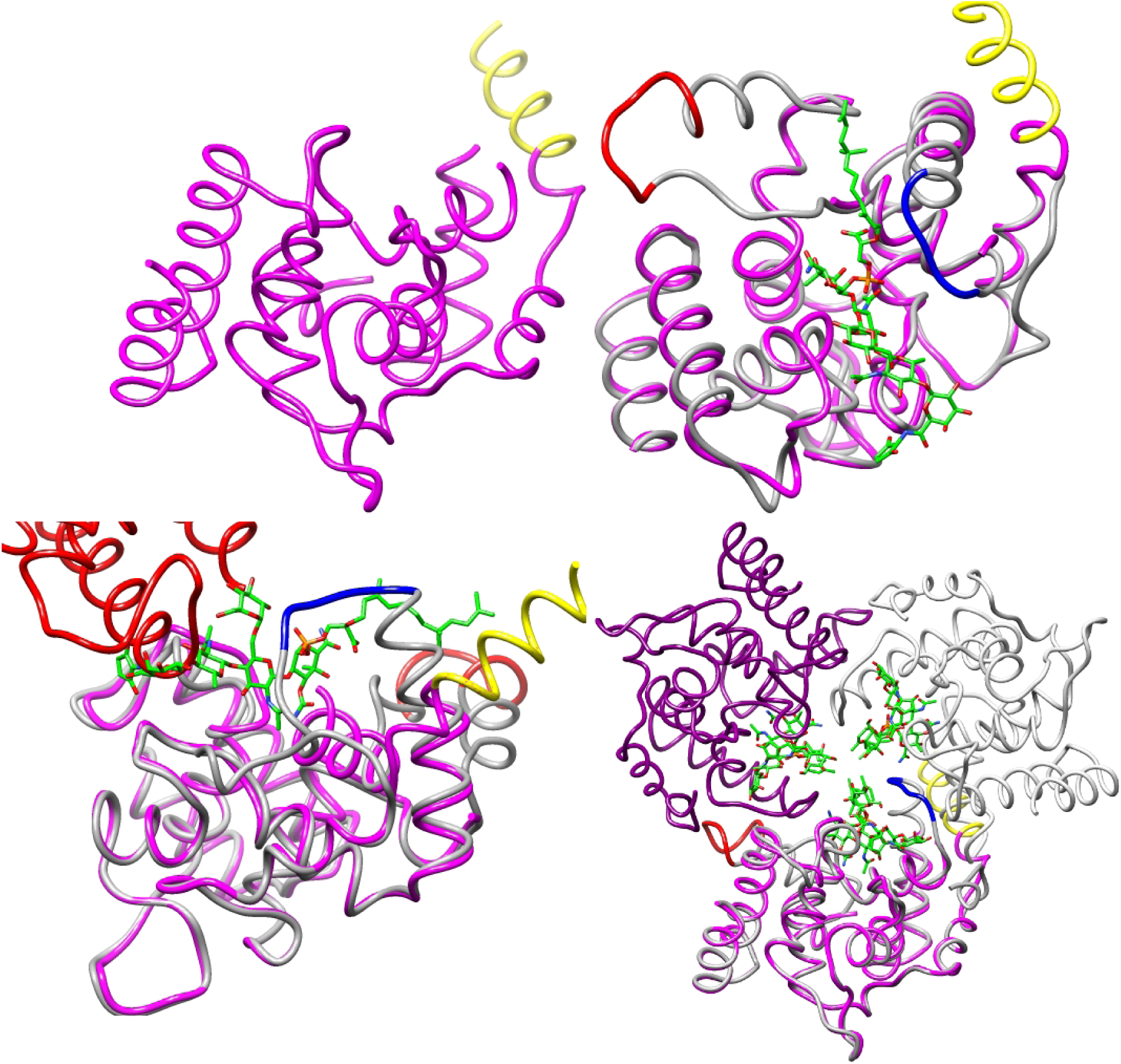
3D structures of 2MGT and 68MGT. A) Top-left. The 3D structure of 2MGT. Residues belonging to the TM-domain (I53-L64) are colored yellow. B) Top-right. The 3D structure of 2MGT superposed onto the 3D structure of 68MGT with bound moeA (green, colored by heteroatom). Element A: apex colored red. Element B: apex colored blue. C) Bottom-left. As B, but also showing a symmetry related 2MGT molecule (red partial chain) that overlaps with the A, B and D-ring (Fig. 1) of the 68MGT bound moeA molecule. This illustrates that the crystallization of 2MGT precludes the binding of moeA. D) Bottom-right. As B, but shown as part of the 68MGT quarternary structure (a trimer with a diameter of ~ 80 Å). The 2MGT TM-domain (yellow) occupies the same coordinates as structural elements of chain C of the 68MGT trimer. Chain A-C are colored dark grey, dark magenta, and light grey, respectively. Figures made with Chimera [44] and GIMP.

**Fig. 3:**
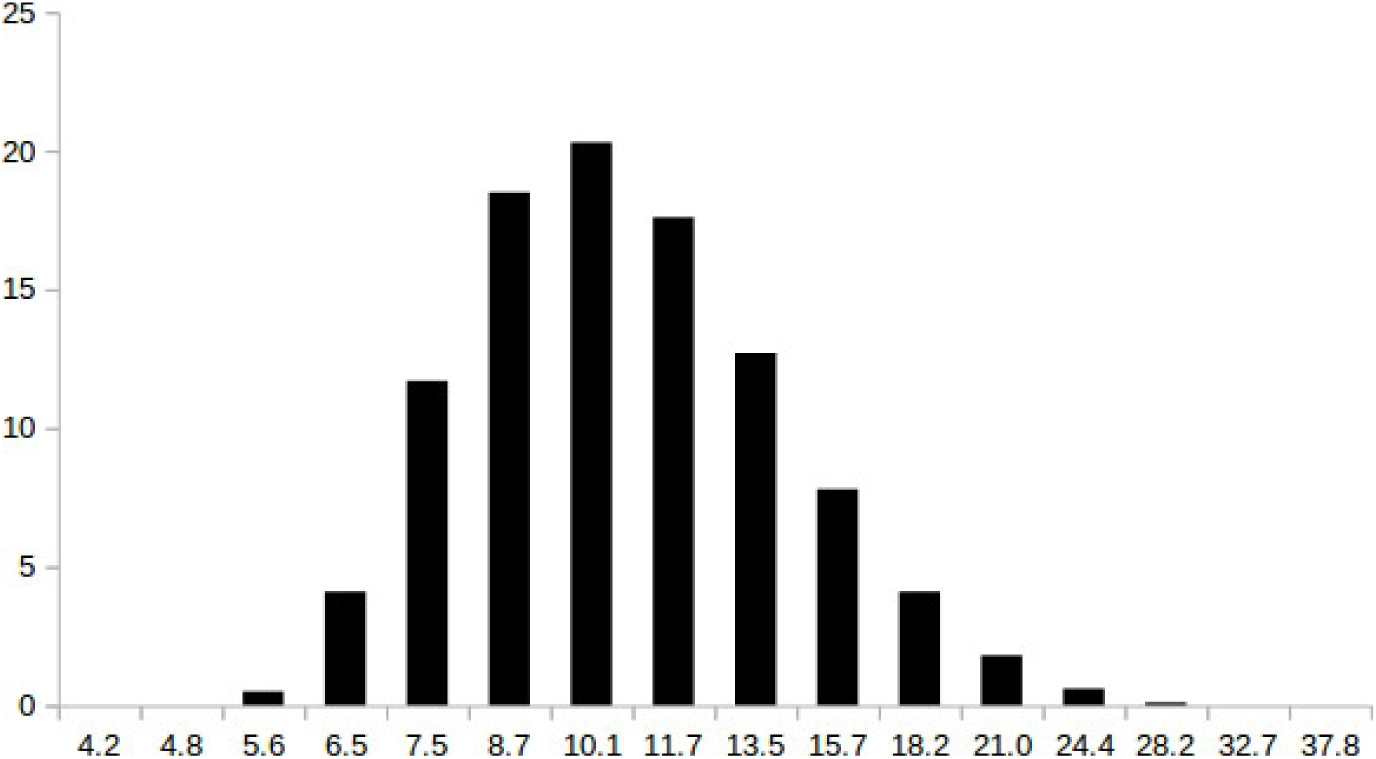
Polydispersity of 6MGT in assay buffer as measured (in duplo) by dynamic light scattering. X-axes: diameter (nm). Y-axis: volume (%). A maximum occurs at 10.1 (11.7) nm at 20.3 (20.3) volume (%) with a Z-average of 17.7 (18.5) nm.

**Fig. 4:**
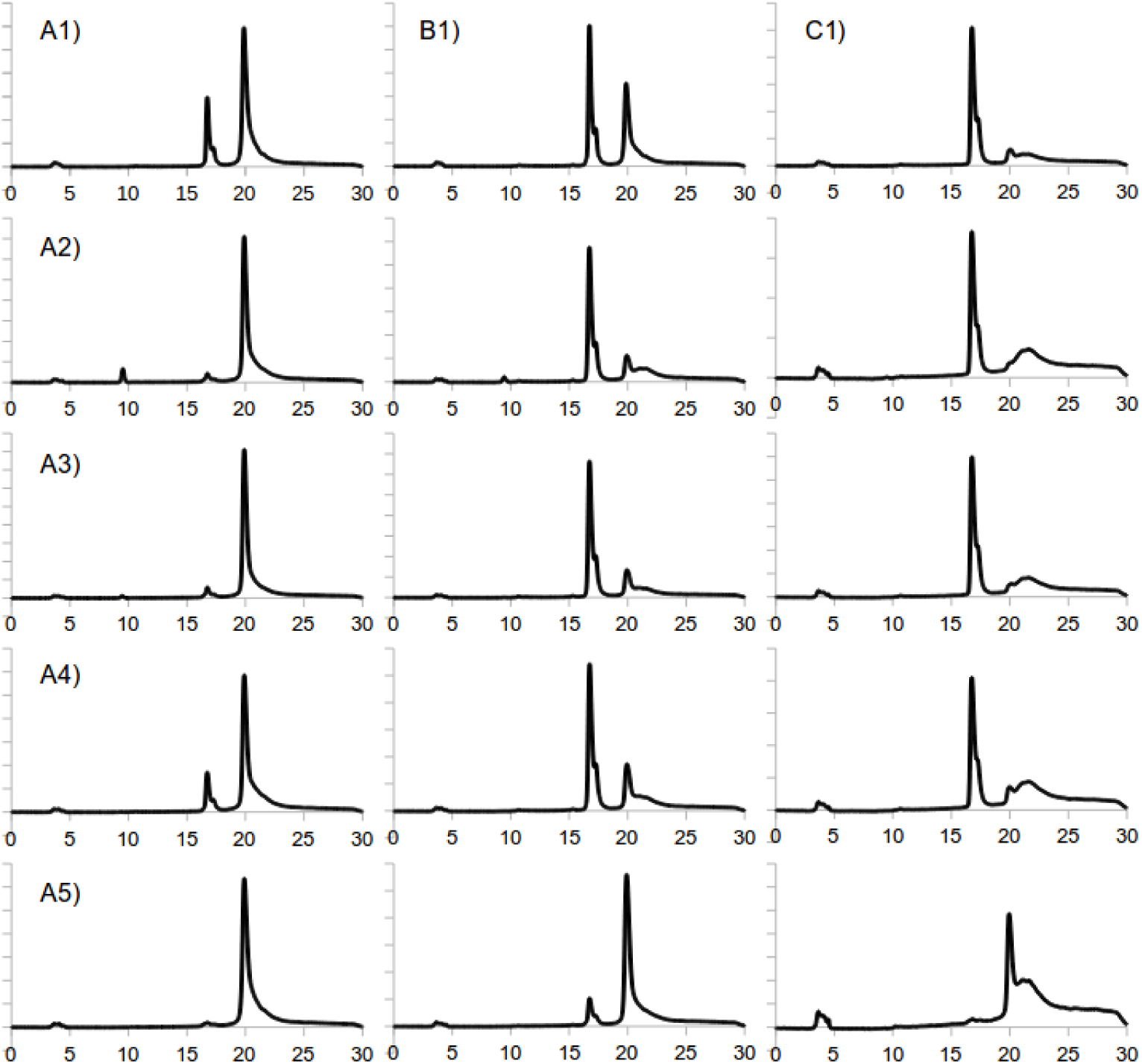
Ion exchange chromatograms showing the conversion of NBD-lipid II (20.1 ml) into NBD-GlcNAc-MurNAc (16.8 ml) by a batch of purified 2MGT and lysozyme under a variety of enzyme assay conditions. A1) 20 mM Na HEPES pH 7.5, 1 mM MnCl_2_ and 1.6 mM C_10_E_8_. A2-4) equals A1, plus either 500, 50, or 5 μM ppb, respectively. A5) equals A1, plus 1 μM moeA. B1-B5) equals A1-A5, but with C_10_E_8_ substituted for 2 mM DM. C1-C5) equals B1-B5, but with DM substituted for 0.01 % v/v *S. aureus* lipids. Assay time: 2 hr (A and B), 10 min (C). X-axis: time (min) with a sampling frequency of 5 Hz. Y-axis: fluorescence units (FLU), with an interval of 100, 50 (C2-C4), or 20 (C5) FLU. Figure made with LibreOffice Calc/Draw and GIMP (as well as figs. 5&6)

**Fig. 5:**
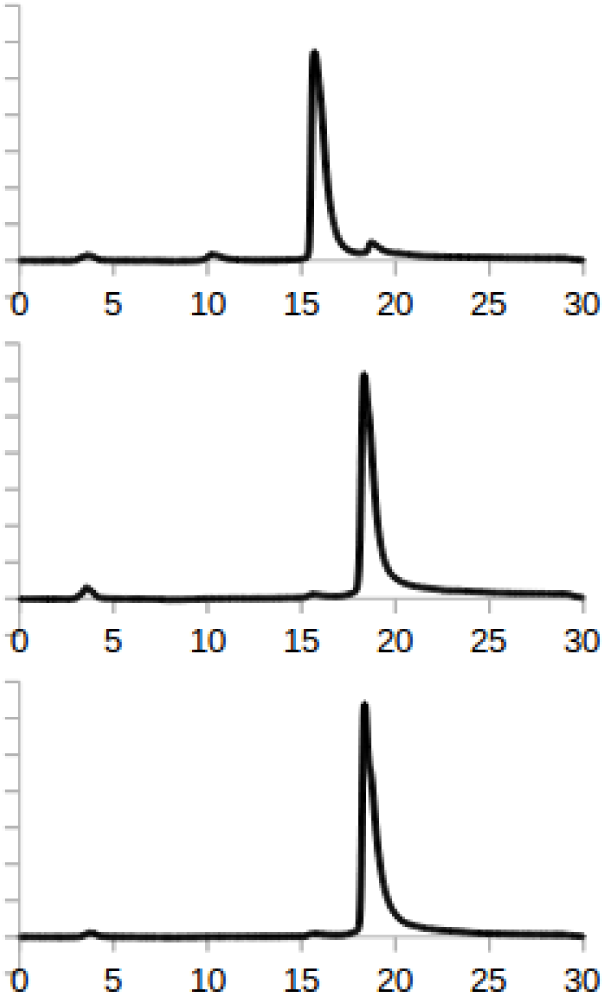
Ion exchange chromatograms showing the conversion of NBD-lipid II (20.1 ml) into NBD-GlcNAc-MurNAc (16.8 ml) by 2MGT and lysozyme in an assay condition containing A) 20 mM Na HEPES pH 7.5, 1 mM MnCl2, 0.1 % v/v *S. aureus* lipids (top). B) As A, plus 16 μM TAN1532B (middle). C) As A, plus 1 μM moeA (bottom). Assay time: 10 min. X-axis: time (min). Y-axis: fluorescence units (FLU). Interval: 100 FLU.

**Fig. 6:**
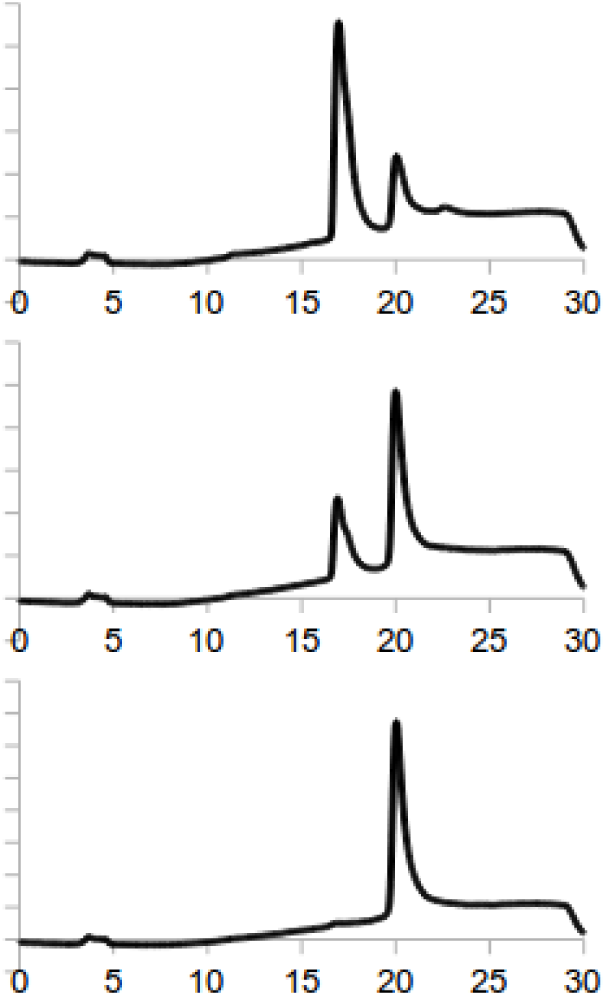
Ion exchange chromatograms showing the conversion of NBD-lipid II (20.1 ml) into NBD-GlcNAc-MurNAc (16.8 ml) by 6MGT and lysozyme in an assay condition containing A) 20 mM Na HEPES pH 7.5, 200 mM NaCl and 1 mM MnCl2 (top). B) As A, plus 5 μM ppb (middle). C) As A, plus 1 μM moeA (bottom). Assay time: 2 hr. X-axis: time (min). Y-axis: fluorescence units (FLU). Interval: 50 FLU.

### Growth of S. aureus cells and extraction of S. aureus lipids – S. aureus

strain ATCC 29213 was used to inoculate 5 ml neutral broth (37 ºC). The cells were grown for 19 h, after which the pre-culture was used to inoculate 2 L neutral broth, and continued for 19 h (37 ºC). The cells were then harvested and frozen (-80 ºC). To 0.9 g *S. aureus* cell pellet (from 200 ml culture) ultra-pure water was added, as well as NaCl (500 mM), lysozyme (100 mg) and DNase I (0.1 mg). The suspension (2.5 ml) was incubated for 2 h at 37 ºC, vortexed every half hour, and transferred into a 100 ml glass vial. While vortexing, 50 ml hexane isopropanol (3:2 [29]) was added to the cell suspension. The mixture was left to settle for 30 min and the solvent harvested. Samples were taken for thin layer chromatography analysis (Fig. S3). The stationary phase was silica gel 60G F254 (Merck), the mobile phase was a mixture of chloroform (65 %), methanol (25 %) and acetic acid (10 %), and staining of the phospholipids was performed with CuSO_4_ (100 mg/ml) [30]. The solvent was subsequently roto-evaporated to dry-ness, which yielded 30 mg *S. aureus* lipid powder. A 2 % wt/vol *S. aureus* lipid solution was subsequently made by adding an appropriate volume of ultra-pure water followed by rigorous vortexing.

### Dynamic light scattering

To investigate the dispersity of size exclusion purified 6MGT, the enzyme was diluted with buffer (20 mM Na HEPES pH 7.5, 1 mM MnCl_2_, 200 mM NaCl) to 0.1 mg/ml, pipetted into a quartz cuvette (QS 3 mm, Hellma) and measured (30 ºC) with a Zetasizer Nano ZS dynamic light scattering system (Fig. 3).

### Modelling

To predict a potential interaction between ppb and the quarternary structure of TM-less 68MGT (Fig. 2D, PDB code 3HZS, a trimer), ppb was drawn in ChemDraw and topologies generated with PRODRG [31] then together with the protein target (without bound moeA, PO_4_, and H_2_O) submitted to SwissDock [32] (Fig. 7).

**Fig. 7:**
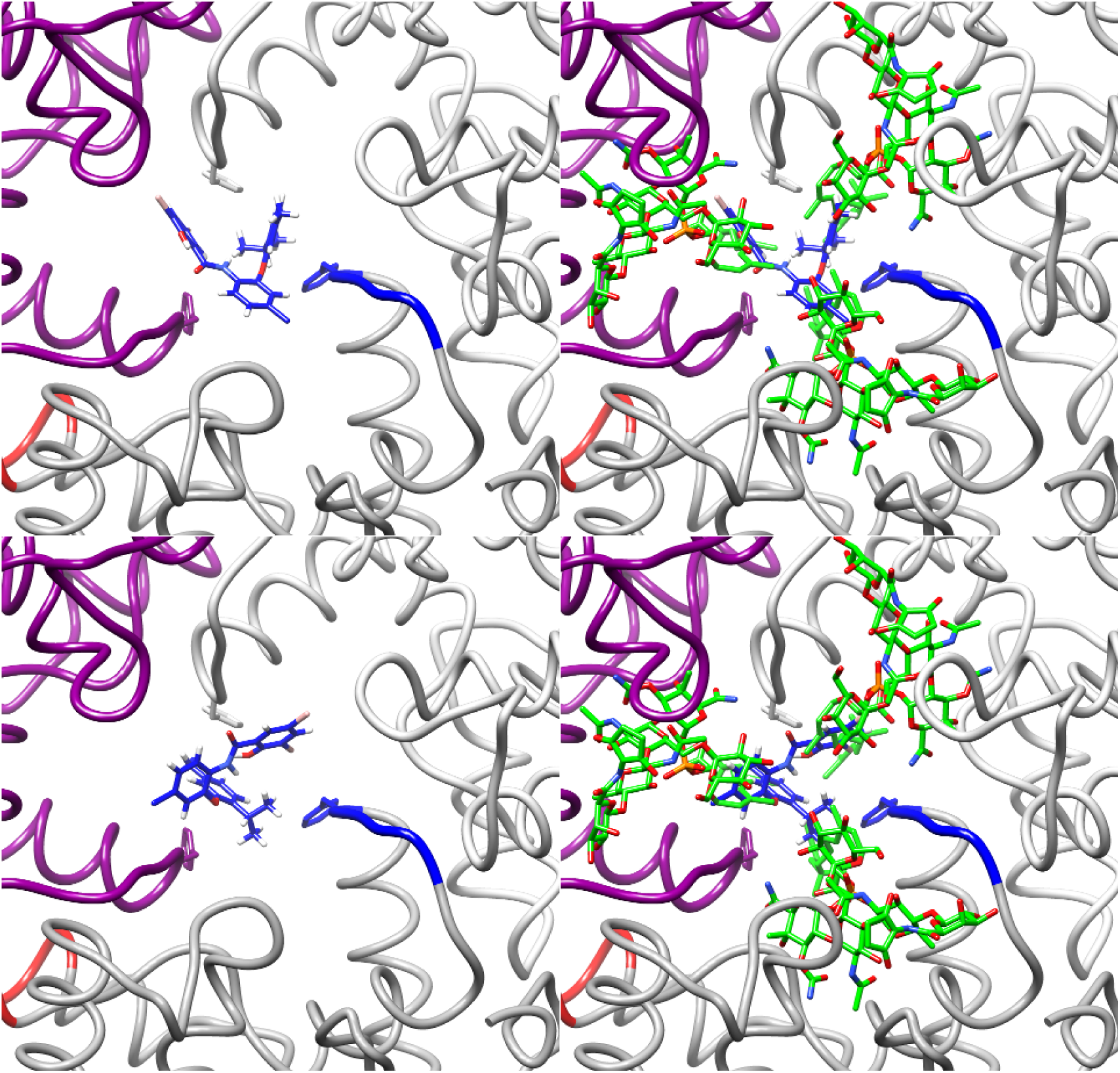
Modelling of the 68MGT (trimer) ppb interaction. A) Top-left. Ppb (blue, colored by heteroatom, ΔG = −7.91) shown bound (via hydrophobic interactions) in an aromatic pocket lined by three F150’s (stick representation) from chains A, B and C (color scheme as in fig. 2D). B) Top-right. As A, but also showing bound moeA. C) Bottom-left. As A, but showing a different model (ΔG = −7.88). D) Bottom-right. As B.

## Results

### SaMGT and SaMGT truncates

In this study we used a monoglycosyltransferase (MGT) from *S. aureus* (*Sa*MGT) to investigate how a non-saccharide salicylanilide (Fig. 1 (**3**), ppb) inhibited the *Sa*MGT-catalyzed lipid II polymerization reaction, as a general model for how the PBP GT-catalyzed reaction was inhibited by salicylanilides. *Sa*MGT is a 31 kDa enzyme (Table 1 (1)) that is composed of three domains: a cytoplasmic domain (residues 1-41), a transmembrane (TM) domain (residues 42-64, ILLKILLTILIIIALFIGIMYFL, fig. 2A), and a membrane-associated periplasmic domain (residues 65-269) that catalyzes a lipid II polymerization reaction. *Sa*MGT degrades during purification (data not shown), unlike Δ28-269 *Sa*MGT (2MGT, table 1 (2)). A 3D structure of 2MGT has been elucidated (PDB code 3VMQ [23], and this research (Table S1, fig. 2A) as well as a 3D structure of a TM-less Δ68-268 *Sa*MGT (68MGT, table 1 (4), PDB code 3HZS, fig. 2B) with bound moenomycin A (moeA)) [33]. 68MGT crystallizes as a trimer (space group H3_2_, fig. 2D), with the interface between chains A, B and C burying ~400 Å^2^ out of a total surface area of ~11000 Å^2^ per chain (PISA [34]), whereas 6MGT (table 1 (3)), a TM-less *Sa*MGT construct analogous to TM-less 68MGT, oligomerizes in solution (Fig. 3), just as TM-less Δ68-268 (table 1 (5)) [35] and TM-less Δ68-269 *Sa*MGT (table 1 (6)) [36].

### Structural investigation into 2MGT ppb inhibition

In an attempt to gain insight into the inhibition of the *Sa*MGT-catalyzed reaction by ppb (Fig. 1 (**3**)), 2MGT was purified as described (for reasons of simplicity) and initially crystallized in the absence of ppb. Several well diffracting protein crystals were obtained that crystallized in an orthorhombic space group. 2MGT crystallizes as a monomer (Table S1) and its 3D structure is similar to the 3D structure of 2MGT deposited under PDB code 3VMQ (2 molecules per ASU) [23] with an RMSD over 180 Cα-atoms of 0.4 Å (chain A of 3VMQ) and 0.7 Å (chain B). The 3D structure is further similar to the 3D structure of moeA bound TM-less 68MGT with an RMSD of 0.8 Å over 161 Cα-atoms (monomer A). The 2MGT 3D structure has three three flexible elements, which are, as a consequence, only partly defined by electron density. One of these elements is the N-terminal α-helix (Fig. 2A). The other two elements are formed by flexible structural elements comprised of amino acids 109 through 132 (element A, hypothesized to be important for glycan processivity [37], fig. 2B) and amino acids 145 through 154 (element B, fig. 2B). However, when attempts were subsequently made to elucidate a 3D structure of 2MGT with bound ppb (after overnight exposure of purified 2MGT to a 50-fold molar excess of ppb (as powder) and co-crystallization of 2MGT with ppb) the elucidated 2MGT 3D structure failed to reveal bound ppb.

### Ppb inhibition of 2MGT

To verify whether ppb inhibited the *Sa*MGT-catalyzed reaction, the enzymatic activity of purified 2MGT was tested using the protocol published by Wu *et al*. [16]. This revealed that 2MGT was active and polymerized NBD-lipid II (lipid II) and that the presence of moeA or ppb in the assay condition led to the inhibition of the 2MGT-catalyzed reaction (data not shown). The enzyme assay conditions were then altered to more closely resemble the assay conditions used by Terrak *et al*. [35]. 2MGT was found to be active and to be inhibited by moeA and ppb (Fig. 4 (A1-A5)). C_10_E_8_ (octaethylene glycol monodecyl ether, decyl-PEG) was subsequently substituted for n-decyl-β-D-maltopyranoside (DM). 2MGT polymerized lipid II, moeA inhibited 2MGT, but ppb did not inhibit the 2MGT-catalyzed lipid II polymerization reaction (Fig. 4 (B1-B5)).

### Isolation of lipids from S. aureus cells and inhibition of 2MGT by ppb

To further investigate this result, C_10_E_8_ and DM were substituted for lipids directly isolated from *S. aureus* cells (Fig. S3). The *S. aureus* extract contained two types of lipids with R_f_ values similar to lipids present in the *E. coli* total lipid extract, and one lipid (nearest to the spot origin, presumably lysyl-phosphatidylglycerol (LPE) [30]) not present in the *E. coli* lipid extract and thus specific to *S. aureus*. 2MGT polymerized the lipid II substrate (Fig. 4 (C1)) and the presence of moeA in the assay condition led to the inhibition of 2MGT (Fig. 4 (C5), Fig. 5C). 2MGT was not inhibited when the enzyme assay condition contained ppb (Fig. 4, C2-C4), but 2MGT was inhibited in the presence of TAN1532B (Fig. S4 (**4**), fig. 5B).

### Inhibition of TM-less 6MGT by ppb and modelling of the 68MGT ppb interaction

A final experiment was subsequently devised and ppb inhibition of TM-less 6MGT tested in an enzyme assay condition (without detergent) in which 6MGT oligomerized (Fig. 3). 6MGT was active and polymerized lipid II (Fig. 6A), was inhibited when the assay condition contained moeA (Fig. 6C), and, surprisingly when the assay condition contained ppb (Fig. 6B). The interaction between ppb and oligomeric 6MGT was modeled with the 3D structure of trimeric 68MGT, which revealed ppb bound, in various orientations (models), in a hydrophobic pocket formed by chain A, B and C of the 68MGT trimer (Fig. 7A, C).

## Discussion

### Structural investigations into glycosyltransferase inhibition by ppb, a non-saccharide salicylanilide inhibitor

In this study we attempted to elucidate a 3D structure of Δ28-269 *Sa*MGT (2MGT) with bound phenylphenoxybenzamide (Fig. 1 (**3**), ppb) with the aim to facilitate structure-based salicylanilide (antibiotic) design. Despite multiple attempts, a 3D structure of 2MGT with bound ppb was eventually not elucidated. To explain this lack of success, it was hypothesized that 2MGT had not bound ppb during ppb treatment and co-crystallization or had bound ppb, but had bound ppb through its flexible TM-domain (hydrophobic) and therefore could not be resolved. Alternatively, it was hypothesized that 2MGT had bound ppb, but could not maintain bound ppb upon crystallization, as exemplified by the inability of crystallized 2MGT to bind moenomycin A (moeA) (Fig. 2C) or as a consequence of crystallization induced non-specific contacts [38], as exemplified by element A (in)flexibility in the 3D structures of 2MGT and Δ28-268 *Sa*MGT (68MGT) (Fig. 2B). Last, it was hypothesized that ppb was not an inhibitor of 2MGT, but appeared to be, as a consequence of the protocol used [16] to assay glycosyltransferase (GT) ppb inhibition.

### Ppb is not a GT inhibitor

To test this hypothesis, we employed a) 2MGT to exclude the possibility that enzyme assay results were influenced by domains found in structurally more complex PBPs (see Introduction) and b) varied enzyme assay conditions to probe whether ppb inhibition of 2MGT enzymatic activity was assay condition-dependent. The protocol outlined by Wu *et al*. [16] was initially used, and the result showed that ppb inhibited 2MGT (data not shown), which suggested that ppb was indeed a GT inhibitor [15]. The enzyme assay condition was then altered to resemble the enzyme assay condition used by Terrak *et al*. [36] to investigate the kinetic properties of TM-less Δ68-268 *Sa*MGT (Table 1 (6)). However, organic solvents were omitted to avoid assay component concentration changes as a consequence of MeOH evaporation, and to avoid the possibility that the solvents affected the 2MGT native structure [39]. The results showed that 2MGT was active, was inhibited by moeA and was inhibited by ppb (Fig. 4 (A1-A5)). This experiment revealed that the inhibition of 2MGT by ppb, despite substantial assay condition changes, could be reproduced. However, upon termination of the 2MGT-catalyzed reaction, and analysis of the reaction mixture contents, a substantial amount of lipid II substrate remained unconverted (Fig. 4 (A1)). This was attributed to the temperature used to perform the assay (30 ºC) and to lipid II substrate aggregation in the absence of organic solvents [40]. Next, C_10_E_8_ was substituted for the disaccharide-based detergent n-decyl-β-D-maltopyranoside (DM) and the experiment repeated. 2MGT was active, 2MGT was inhibited by moeA, but 2MGT was not inhibited by ppb (Fig. 4 (B1-B5)). These results suggested that ppb was not a GT inhibitor, contrary to earlier observations, and that ppb inhibition of 2MGT was detergent-dependent. For this reason, C_10_E_8_, DM and artificial detergents in general, were dispensed with, and substituted for lipids directly extracted from *S. aureus* cells (Fig. S3) via a method previously described by Hara *et al*. [40] and validated by Kolarovic *et al*. [41] with the aim to mimic the native lipid environment of *Sa*MGT, the bacterial plasma membrane. This allowed the enzyme assay time to be reduced from ≥ 2 hr (C_10_E_8_ and DM) to ≤ 10 min (Fig. 4 (C1)). A similar observation was made by Newman *et al*. [42], who showed that lactose permease enzymatic activity increased significantly when reconstituted into liposomes prepared from crude *E. coli* phospholipid extract instead of octyl-β-D-glucopyranoside. Under these enzyme assay conditions, 2MGT was inhibited by moeA (Fig. 4 (C5), Fig. 5C), but ppb failed to inhibit the 2MGT-catalyzed reaction (Fig. 4 (C2-C4)). Thus while moeA inhibited 2MGT effectively under all assay conditions in comparison to ppb, ppb inhibition of 2MGT was C_10_E_8_-dependent, which showed that ppb was not a genuine GT inhibitor, a conclusion further strengthened by the inhibition of 2MGT by TAN1532B (Fig. 5B) [16].

### Ppb binds 6MGT and modelling of the 68MGT ppb interaction

To further validate this conclusion, 2MGT was substituted for TM-less 6MGT, which oligomerized under assay conditions without detergent (Fig. 3) in accordance with TM-less Δ68-269 [35] and Δ68-268 *Sa*MGT [36]. 6MGT was active and polymerized lipid II, moeA inhibited 6MGT and, contrary to expectations, 6MGT was inhibited by ppb (Fig. 6B). However, inspection of the 3D structure of 2MGT revealed that it’s periplasmic domain (6MGT) lacked a pocket lined with hydrophobic residues that could accommodate ppb. It was therefore hypothesized that ppb bound/sequestered lipid II. This was proven incorrect after Wu *et al*. [16] showed that TM-less and TP-less truncates of *A. baumannii* PBP1b and *C. difficile* PBP bound ppb in the absence of lipid II. The fact that 2MGT crystallized as a monomer, whereas TM-less Δ68-268 *Sa*MGT (68MGT) crystallized as a trimer (Fig. 2D) and TM-less 6MGT, under assay conditions, assembled into oligomers, among which trimers (Figs. 2D & 3) led to the hypothesis that ppb bound oligomeric rather than monomeric 6MGT. To investigate this hypothesis, a docking experiment was performed with 68MGT (trimer) and ppb, as a model for how a trimer of 6MGT may bind ppb. This experiment revealed that ppb bound a hydrophobic pocket lined by the F150’s of chain A, B and C of 68MGT (I186 and L227 in *Cd*PBP and *Ab*PBP1b, respectively) and showed that bound ppb simultaneously occupied the moenocinol side chain binding pockets of moeA (Fig. 7B, D), which explained why ppb, under assay conditions, inhibited the 6MGT-catalyzed lipid II polymerization reaction. However, oligomerization of Δ68-269 and Δ68-268 *Sa*MGT was shown to be detergent-sensitive [35, 36], and comparison of the 3D structure of 68MGT with that of 2MGT (Fig. 2D) showed that steric hindrance would prevent 2MGT monomers from assembling into the same configuration as 68MGT monomers upon trimerization. This suggested that 2MGT would not be able to bind ppb even in an oligomerization event, in contrast to 68MGT. For these reasons, ppb inhibition of 6MGT, under assay conditions, was concluded to represent an artifact. Our investigations thus reveal that construct choice, and detergent choice strongly affect GT assay results and show that it is imperative that GT assays are thoroughly validated before being applied to screen for GT inhibitors, in order to avoid false-positives [43].

## Acknowledgements

We thank the technical services provided by the “Synchrotron Radiation Protein Crystallography Facility of the National Core Facility Program for Biotechnology, Ministry of Science and Technology” and the “National Synchrotron Radiation Research Center”, a national user facility supported by the Ministry of Science and Technology of Taiwan, ROC. In addition, GGW would like to thank Guo, Chih-Wei for help with analyzing Fig. S3.

## References

1: Taubes G. – The bacteria fight back – Science, 2008, 321(5887), 356–361

2: Perkins H.R. – Chemical structure and biosynthesis of bacterial cell walls – Bacteriol. Rev., 1963, 27(1), 18–55

3: Anderson J.S., Matsuhashi M., Haskin M.A., Strominger J.L. – Lipid-phosphoacetylmuramylpentapeptide and lipid-phosphodisaccharide-pentapeptide: presumed membrane transport intermediates in cell wall synthesis – Proc. Natl. Acad. Sci. U.S.A., 1965, 53(4), 881–889

4: Higashi Y., Strominger J.L., Sweeley C.C. – Biosynthesis of the peptidoglycan of bacterial cell walls. XXI. Isolation of free C55-isoprenoid alcohol and of lipid intermediates in peptidoglycan synthesis from *Staphylococcus aureus* – J. Biol. Chem., 1970, 245(14), 3697–3702

5: Sauvage S., Kerff F., Terrak M., Ayala J.A., Charlier P. – The penicillin-binding proteins: structure and role in peptidoglycan synthesis – FEMS Microbiol. Rev., 2008, 32(2), 234–258

6: Perlstein D.L., Zhang Y., Wang T.S., Kahne D.E., Walker S. – The direction of glycan chain elongation by peptidoglycan glycosyltransferases – J. Am. Chem. Soc., 2007, 129(42), 12674–12675

7: Lovering A.L., de Castro L.H., Lim D., Strynadka N.C.J. – Structural insight into the transglycosylation step of bacterial cell-wall synthesis – Science, 2007, 315(5817), 1402–1405

8: Scheurwater E., Reid C.W., Clarke A.J. – Lytic transglycosylases: bacterial space-making autolysins – Int. J. Biochem. Cell Biol., 2008, 40(4), 586–591

9: King D.T., Strynadka N.C.J. – The mechanism of resistance to β-lactam antibiotics. In: Gotte M., Sheppard D. (eds) Handbook of antimicrobial resistance, 2017, 177–201, Springer, New York, NY

10: Kong K-F., Schneper, L., Mathee K. – Beta-lactam antibiotics: from antibiosis to resistance and bacteriology – APMIS, 2010, 118(1), 1–36

11: Ostash B., Walker S. – Moenomycin family antibiotics: chemical synthesis, biosynthesis, biological activity – Nat. Prod. Rep., 2010, 27(11), 1594–1617

12: Zuegg J., Muldoon C., Adamson G., McKeveney D., Le Thanh G., Premraj R., Becker B., Cheng M., Elliot A.G., Huang J.X., Butler M.S., Bajaj M., Seifert J., Singh L., Galley N.F., Roper D.I., Lloyd A.J., Downson C.G., Cheng T.J., Cheng W.C., Demon D., Meyer E., Meutermans W., Cooper, M.A. – Carbohydrate scaffolds as glycosyltransferase inhibitors with *in vivo* antibacterial activity – Nat. Commun., 2015, doi: 10.1038/ncomms8719

13: Dumbre S., Derouaux A., Lescrinier E., Piette A., Joris B., Terrak M., Herdewijn, P. – Synthesis of modified peptidoglycan precursor analogues for the inhibition of glycosyltransferases – J. Am. Chem. Soc., 2012, 134(22), 9343–9351

14: Derouaux A., Sauvage E., Terrak M. – Peptidoglycan glycosyltransferase substrate mimics as templates for the design of new antibacterial drugs – Front. Immunol., 2013, doi: 10.3389/fimmu.2013.00078

15: Cheng T-J., Wu Y-T, Yang S-T, Lo K-H, Chen S-K, Chen Y-H, Huang W-I, Yuan C-H, Guo, CW, Huang L-Y, Chen K-T, Shih H-W, Cheng Y-S, Cheng W-C, Wong C-H. – High-throughput identification of antibacterials against methicillin-resistant *Staphylococcus areus* (MRSA) and the transglycosylase – Bioorg. Med. Chem., 2010, 18(24), 8512–8529

16: Wu W-S., Cheng W-C, Cheng T-R, Wong C-H. – Affinity-based screen for inhibitors of bacterial transglycosylase – J. Am. Chem. Soc., 2018, 140(8), 2752–2755

17: Bradford M.M. – A rapid and sensitive method for the quantitation of microgram quantities of protein utilizing the principle of protein-dye binding – Anal. Biochem., 1976, 72(1-2), 248–254

18: Ujwal R., Bowie J.U. – Crystallizing membrane proteins using lipidic bicelles – Methods, 2011, 55(4), 337–341

19: Battye T.G.G., Kontogiannis L., Johnson O., Powel H.R., Leslie, A.G.W. – iMOSFLM: a new graphical interface for diffraction-image processing with MOSFLM – Acta Crystallogr. D Biol. Crystallogr., 2011, 67(4), 271–281

20: Strong M., Sawaya M.R., Wang S., Phillips M., Cascio D., Eisenberg D. – Toward the structural genomics of complexes: Crystal structure of PE/PPE protein complex from *Mycobacterium tuberculosis* – Proc. Natl. Acad. Sci. U.S.A., 2006, 103(21), 8060–8065

21: Matthews B.W. – Solvent content of protein crystals – J. Mol. Biol., 1968, 33(2), 491–497

22: McCoy A.J., Grosse-Kunstleve R.W., Adams P.D., Winn M.D., Storoni L.C., Read R.J. – Phaser crystallographic software – J. Appl. Cryst., 2007, doi: 10.1107/S0021889807021206

23: Huang C-Y., Shih H.W., Lin L-Y, Tien Y-W, Cheng T-J, Cheng W-C, Wong C-H, Ma C. – Crystal structure of *Staphylococcus areus* transglycosylase in complex with a lipid II analog and elucidation of peptidoglycan synthesis mechanism – Proc. Natl. Acad. Sci. U.S.A., 2012, 109(17), 6496–6501

24: Murshudov G.N., Vagin A.A., Dodson E.J. – Refinement of macromolecular structures by maximum-likelihood method – Acta Crystallogr. D Biol. Crystallogr., 1997, doi: 10.1107/S0907444996012255

25: Emsley P., Lohkamp B., Scott W.G., Cowtan K., – Features and development of Coot – Acta Crystallogr. D Biol. Crystallogr., 2010, doi: 10.1107/S0907444910007493

26: Winn M.D., Ballard C.C., Cowtan K.D., Dodson E.J., Emsley P., Evans P.R., Keegan R.M., Krissinel E.B., Leslie A.G., McCoy A., McNicholas S.J., Murshudov G.N., Pannu N.S., Potterton E.A., Powell H.R., Read R.J., Vagin A., Wilson K.S. – Overview of the CCP4 suite and current developments – Acta Crystallogr. D Biol. Crystallogr., 2011, doi: 10.1107/S0907444910045749

27: Kall L., Krogh A., Sonnhammer E.L. – A combined transmembrane topology and signal peptide prediction method – J. Mol. Biol., 2004, 338(5), 1027–1036

28: Kurz M., Guba W., Vertesy L. – The three-dimensional structure of moenomycin A, a potent inhibitor of penicillin-binding protein 1b – Eur. J. Biochem., 1998, 252(3), 500–507

29: Hara A., Radin N.S. – Lipid extraction of tissues with a low-toxicity solvent – Anal. Biochem., 1978, 90(1), 420–426

30: Tsai M., Ohniwa R.L., Kato Y., Takeshita S.L., Ohta T., Saito S., Hayashi H., Morikawa K. – *Staphylococcus aureus* requires cardiolipin for survival under conditions of high salinity – BMC Microbiol., 2011, doi: 10.1186/1471-2180-11-13

31: Schuttelkopf A.W., van Aalten D.M. – PRODRG: a tool for high-throughput crystallography of protein-ligand complexes – Acta Crystallogr. D Biol. Crystallogr., 2004, doi: 10.1107/S0907444904011679

32: Grosdidie, A., Zoete V., Michielin O. – SwissDock, a protein-small molecule docking web service based on EADock DSS – Nucleic Acids Res., 2011, doi: 10.1093/nar/gkr366

33: Heaslet H., Shaw B., Mistry A., Miller A.A. – Characterization of the active site of *S. aureus* monofunctional glycosyltransferase (Mtg) by site-directed mutation and structural analysis of the protein complexed with moenomycin – J. Struct. Biol., 2009 167(2), 129–135

34: Krissinel E., Henrick K. – Inference of macromolecular assemblies from crystalline state – J. Mol. Biol., 2007, 372(3), 774–797

35: Wang Q.M., Peery R.B., Johnson R.B., Alborn W.E., Yeh W.K., Skatrud P.L. – Identification and characterization of a monofunctional glycosyltransferase from *Staphylococcus aureus* – J. Bacteriol., 2001, 183(16), 4779–4785

36: Terrak M., Nguyen-Disteche M. – Kinetic characterization of the monofunctional glycosyltransferase from *Staphylococcus aureus* – J. Bacteriol., 2006, 188(7), 2528–2532

37: Yuan Y., Barrett D., Zhang Y., Kahne D. Sliz P., Walker S. – Crystal structure of a peptidoglycan glycosyltransferase suggests a model for processive glycan chain synthesis – Proc. Natl. Acad. Sci. U.S.A., 2007, 104(13), 5348–5353

38: Luo J., Liu Z., Guo Y., Li M. – A structural dissection of large protein-protein crystal packing contacts – Sci. Rep., 2015, doi: 10.1038/srep14214

39: Bisswanger H. – Enzyme assays – Pers. in Sci., 2014, doi: 10.1016/j.pisc.2014.02.005

40: Feng B.Y., Shoichet, B.K. – A detergent-based assay for the detection of promiscuous inhibitors – Nat. Protoc., 2006, 1(2), 550–553

41: Kolarovic L., Fournier N.C. – A comparison of extraction methods for the isolation of phospholipids from biological sources – Anal. Biochem., 1986, 156(1), 244–250

42: Newman M.J., Wilson T.H. – Solubilization and reconstitution of the lactose transport system from *Escherichia coli* – J. Biol. Chem., 1980, 255(22), 10583–10586

43: Baell J., Walters M.A. – Chemical con artists foil drug discovery – Nature, 2014, 513(7519), 481–483

44: Pettersen E.F., Goddard T.D., Huang C.C., Couch G.S., Greenblatt D.M., Meng E.C., Ferrin T.E. – UCSF Chimera–A visualization system for exploratory research and analysis – J. Comput. Chem., 2004, 25(13), 2004, 1605–1612

